# FIREBALL: A tool to fit protein phase diagrams based on mean-field theories for polymer solutions

**DOI:** 10.1101/2023.03.19.533322

**Authors:** Mina Farag, Alex S. Holehouse, Xiangze Zeng, Rohit V. Pappu

## Abstract

Biomolecular condensates form via phase transitions of condensate-specific biomacromolecules. Intrinsically disordered regions (IDRs) featuring the appropriate sequence grammar can contribute homotypic and heterotypic interactions to the driving forces for phase separation of multivalent proteins. At this juncture, experiments and computations have matured to the point where the concentrations of coexisting dense and dilute phases can be quantified for individual IDRs in complex milieus both *in vitro* and *in vivo*. For a macromolecule such as a disordered protein in a solvent, the locus of points that connects concentrations of the two coexisting phases defines a phase boundary or binodal. Often, only a few points along the binodal, especially in the dense phase, are accessible for measurement. In such cases and for quantitative and comparative analysis of parameters that describe the driving forces for phase separation, it is useful to fit measured or computed binodals to well-known mean-field free energies for polymer solutions. Unfortunately, the non-linearity of the underlying free energy functions makes it challenging to put mean-field theories into practice. Here, we present FIREBALL, a suite of computational tools designed to enable efficient construction, analysis, and fitting to experimental or computed data of binodals. We show that depending on the theory being used, one can also extract information regarding coil-to-globule transitions of individual macromolecules. Here, we emphasize the ease-of-use and utility of FIREBALL using examples based on data for two different IDRs.

**Statement of Significance:** Macromolecular phase separation drives the assembly of membraneless bodies known as biomolecular condensates. Measurements and computer simulations can now be brought to bear to quantify how the concentrations of macromolecules in coexisting dilute and dense phases vary with changes to solution conditions. These mappings can be fit to analytical expressions for free energies of solution to extract information regarding parameters that enable comparative assessments of the balance of macromolecule-solvent interactions across different systems. However, the underlying free energies are non-linear and fitting them to actual data is non-trivial. To enable comparative numerical analyses, we introduce FIREBALL, a user-friendly suite of computational tools that allows one to generate, analyze, and fit phase diagrams and coil-to-globule transitions using well-known theories.

## Main Text

Biomolecular condensates are membraneless bodies that confer spatiotemporal control over various cellular processes (1–4), including ribosomal assembly (5,6), transcription (7–9), autophagy (10–13), stress response (14–17), and cellular trafficking (18–20). Condensates form via coupled associative and segregative phase transitions of multivalent biomacromolecules, namely protein and RNA molecules (21). Although intrinsically disordered proteins (IDPs) are not required for phase separation, in many cases they have been found to be major drivers (22).

Phase diagrams offer a clear way to quantify and compare the phase behaviors of macromolecules (23). In many cases, phase diagrams are interpreted using free energy-based models of phase separation, such as Flory-Huggins theory (FHT) (24–26). Although IDPs are not homopolymers, one can fit the FHT model to measured data for binodals and do so with high accuracy. Indeed, achieving good fits to measured and computed binodals helped uncover the insight that prion-like low-complexity domains behave like effective homopolymers (27). This is due to conserved sequence patterning whereby stickers, which are the cohesive motifs, are uniformly and non-randomly distributed along the linear sequence. Therefore, fitting measured or computed binodals to free energies for homopolymers, block copolymers, or associative systems can help identify the class of physical systems to which an IDP belongs. Further, such numerical analysis allows for comparative assessments of sequence-or composition-specific driving forces for phase separation.

Unfortunately, expressions for free energies of polymer solutions contain many non-linear terms, making them analytically intractable. This makes it challenging to deploy these models in numerical analyses of measured or computed binodals. While there has been some work to approximate analytical solutions and provide tools to generate phase diagrams (28–30), we see and perceive the lack of a useful and comprehensive toolkit that allows one to deploy different free energy models in a flexible manner.

The FHT model, which treats polymers as a gas of residues or Kuhn units (21), is one of several models for describing the free energies of polymer solutions. The simplest theories of complex coacervation add a mean-field, Debye-Hückel-like term to capture the effects of screened electrostatic interactions among charged polymers (31). Muthukumar added the contributions of three-body interactions and terms that allow for describing the effects of concentration fluctuations and correlation lengths (32,33). Tanaka’s approaches led to the introduction of contributions from specific molecular associations, acknowledging the presence of stickers or cohesive motifs that enable reversible crosslinking (34–36). Semenov and Rubinstein took a similar approach in their work, and this led to a clarification of the distinctions between associative percolation transitions versus segregative phase separation (37). Lin and Chan developed a generalization of FHT, building on the work of Olvera de la Cruz and colleagues, to include the effects of sequence-specific stickers and patterning effects (38–40). All these theories rest on the core structure of the original FHT, while introducing various additions or generalizations.

A drawback of the FHT model is that it does not account for conformational preferences of polymers in dilute solutions or within dense phases. Conformational considerations are of significant import in dilute solutions, at interfaces, and within dense phases because these contribute to the biochemical reactions influenced by the drivers of phase separation (41). The deficiencies of FHT have been remedied, in part, by the Gaussian Cluster Theory (GCT) of Raos and Allegra (42,43). The use of GCT also paves the way for a theory-based computation of phase diagrams by extracting the two- and three-body interaction coefficients from simulations or measurements of coil-to-globule transitions. This is helpful and relevant because several groups have shown a clear connection between the interactions that determine conformations of individual molecules in dilute phases and the driving forces for phase separation (27,42–50,41). For example, *stickers* that drive phase separation via intermolecular interactions often drive chain compaction in the single-chain limit via intramolecular interactions. In the context of IDPs, *stickers* can be blocks of charged residues or aromatic residues (51,27). Much work has been done to understand the coupling between determinants of single chain conformations in dilute phases and the driving forces for phase separation, and when / how this coupling breaks down. However, being able to connect simulations and experiments to treatments such as the GCT remain challenging. This presents a barrier to quantitative analyses that can be deployed without developing the required theoretical expertise.

We seek to enable easy-to-use numerical analysis of measured or computed binodals. For this, we introduce FIREBALL (FItting REal BinodALs with non-Linear models), a computational tool to fit free energy-based theoretical models to measured or computed binodals. We demonstrate the usage by fitting the FHT model with three-body interactions and the GCT model to measured data. Using the parameters that one obtains from numerical fitting of binodals to analytical theories, one can also construct full phase diagrams, which include binodals, spinodals, and estimates of critical parameters. Additionally, if one fits data to the GCT, then one can obtain quantitative descriptions of coil-to-globule transitions, a topic of considerable interest in the chromatin field.

### Overview of FIREBALL

FIREBALL is a user-friendly computational suite that is accessible via command line tools. It allows one to analyze computed or measured binodals and coil-to-globule transitions using free energy-based theoretical models. FIREBALL is written entirely in Python and is freely available to download via the Pappu Lab GitHub page (https://github.com/pappulab) as an open-source package. FIREBALL also includes documentation that describes the set-up procedure and includes advice for troubleshooting.

#### Methodology for fitting measured or computed binodals

For mean-field free energy-based models of phase separation such as FHT and GCT, concentrations of coexisting phases that define binodals are determined by equalizing the chemical potentials and osmotic pressures across the phase boundary. In practice, this requires drawing a common tangent line between two points on the free energy versus macromolecular concentration curve. The slope of this line is determined by the equalized chemical potential, whereas the intercept along the ordinate, which usually is temperature or a measure of changes to solution conditions, is determined by the equalized osmotic pressure. The two points along the free energy curve that share a common tangent represent the concentrations of coexisting dilute and dense phases. Most theoretical models, including FHT and GCT, contain numerous non-linear terms, making the functional forms analytically intractable. For this reason, one cannot directly calculate the dilute and dense phase concentrations. In contrast, points along the spinodal can be determined by setting the second derivative of the free energy with respect to concentration equal to zero. We use this to our advantage to set up an optimization procedure and determine binodals for a given set of parameters. This optimization uses calculated spinodals, upper and / or lower bounds on concentrations along the binodal given prior measurements of the concentrations at different temperatures, and any system-specific limits on parameters that serve as heuristic bounds for points that make up binodals.

#### Theoretical models included in FIREBALL

Currently, FIREBALL includes support for FHT and GCT. The FHT implementation uses the following functional form for the free energy of a polymer solution:

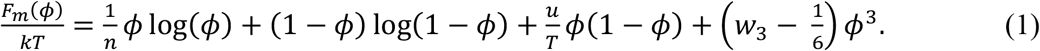

This version of the FHT has been introduced previously (27,52). Here, *ϕ*is the protein volume fraction and *T* is the temperature. *n, u*, and *w*_3_are independent parameters corresponding to the protein length, the strength of two-body interactions, and the strength of three-body interactions. *u*, and *w*_3_are related to the second and third virial coefficients of protein molecules, respectively. The functional form used for the GCT implementation is described in detail by Zeng et al. (53). Treating the system as an effective homopolymer, the GCT model includes θ, *w*_2_, *w*_3_, and *n* as independent parameters that correspond to the theta temperature, the two-body interaction coefficient, the three-body interaction coefficient, and the polymer length. This model also requires an approximate Flory characteristic ratio, Kuhn length, and an average length for the repeating unit. While both models are non-linear, the GCT formalism includes an added complication in the form of *q*, the single-chain compression factor, which is only defined implicitly as a function of *ϕ*and *T*. Thus, we must perform an interpolation for *q* at every distinct temperature to estimate an explicit form for *q* as a function of ϕ. As a result, phase diagram fits performed using FHT require ∼5 seconds to perform, whereas fits performed using GCT require a few minutes. In contrast, fits to measured or computed coil-to-globule transitions using GCT require ∼5 seconds.

### Analyses included within FIREBALL

FIREBALL includes four main functions that can be run directly via the command line. Arguments for all functions are explicitly defined in the FIREBALL documentation.

One can use *fireball-fit* to fit a binodal to a dataset using a specified free energy model. The user may choose initial guesses for parameters, as well as which parameters are “fixed” vs. “free.” Upon running *fireball-fit*, the program will generate a binodal based on the given parameters, calculate the error between the generated binodal and the given dataset, modulate the free parameters, and re-generate a binodal. These steps are repeated until the calculated error is below a given value or the number of cycles exceeds a given threshold. Because concentrations along binodals, especially low-concentration arms of binodals, vary by orders of magnitude, errors are calculated on a logarithmic scale. By default, this optimization employs a Nelder-Mead algorithm, a gradient-free method for determining local minima (54). However, this method can be altered as desired. *fireball-fit* then saves the fitted binodal, spinodal, and fitting parameters and plots the binodal alongside the data. Examples of this output are shown in **Fig. 1** using an FHT-based fit and in **Fig. 2** using a GCT-based fit.

**Fig. 1:**
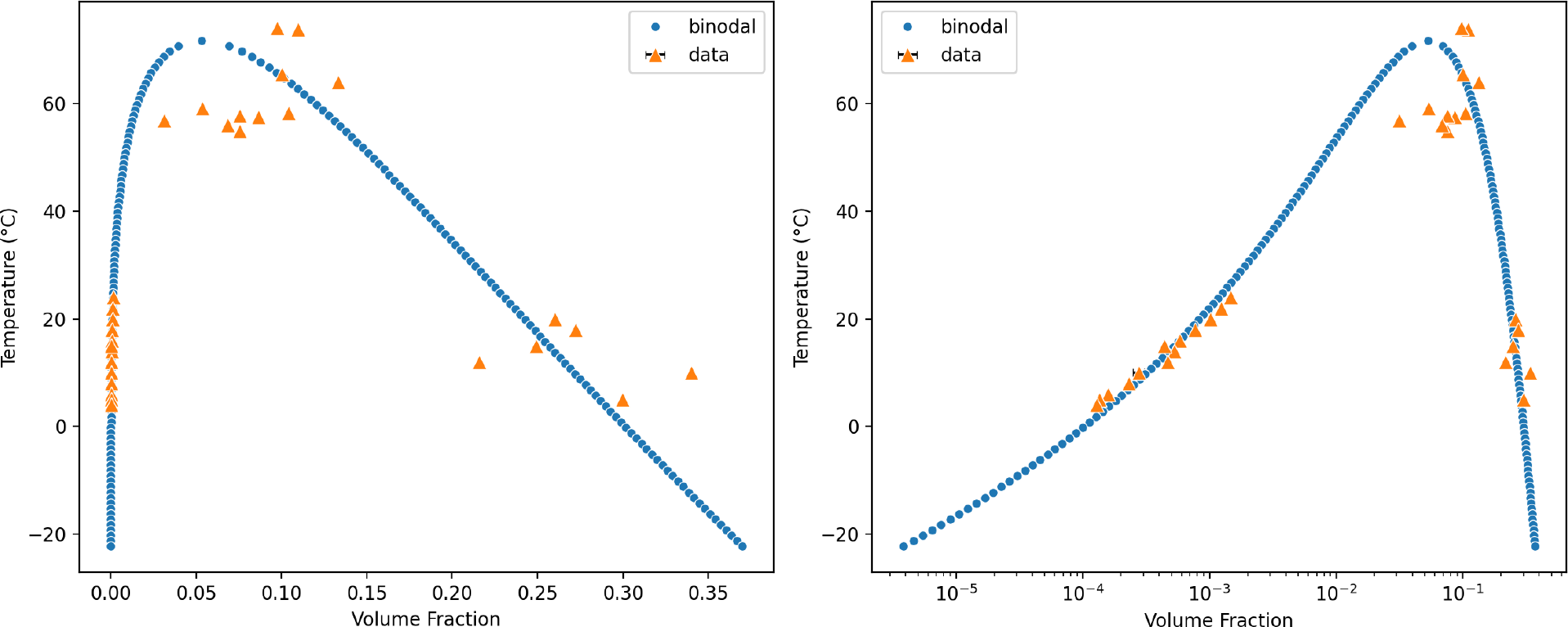
Example output from *fireball-fit* applied to phase separation data of the wild-type A1-LCD from Bremer et al. (48) **using FHT**. This is a two-parameter fit using FHT where *n* is fixed at 137, the length of the A1-LCD protein, and *u* and *w*_3_ are free parameters. Protein concentration is converted to volume fraction by assuming that a volume fraction of 1.0 corresponds to a mass concentration of 1310 mg/mL. The final fitting parameters are *u* = 219 K and *w*_3_ = 0.413. The right panel is identical to the left one, save that it uses a logarithmic scale for the abscissa. This is important because while the measured estimates of dense phase concentrations are noisy, mostly due to pipetting errors, they vary minimally. However, and in contrast to what most computer simulations show (47, 49, 50), the measured dilute phase concentrations vary by 4-5 orders of magnitude, being very low at low temperatures, and approaching the dense phase concentrations as the critical point is approached. The critical point is defined by large fluctuations and the accompanying critical opalescence makes it difficult to estimate using experiments. The result is significant noise as can be seen in the data around the critical point.

**Fig. 2:**
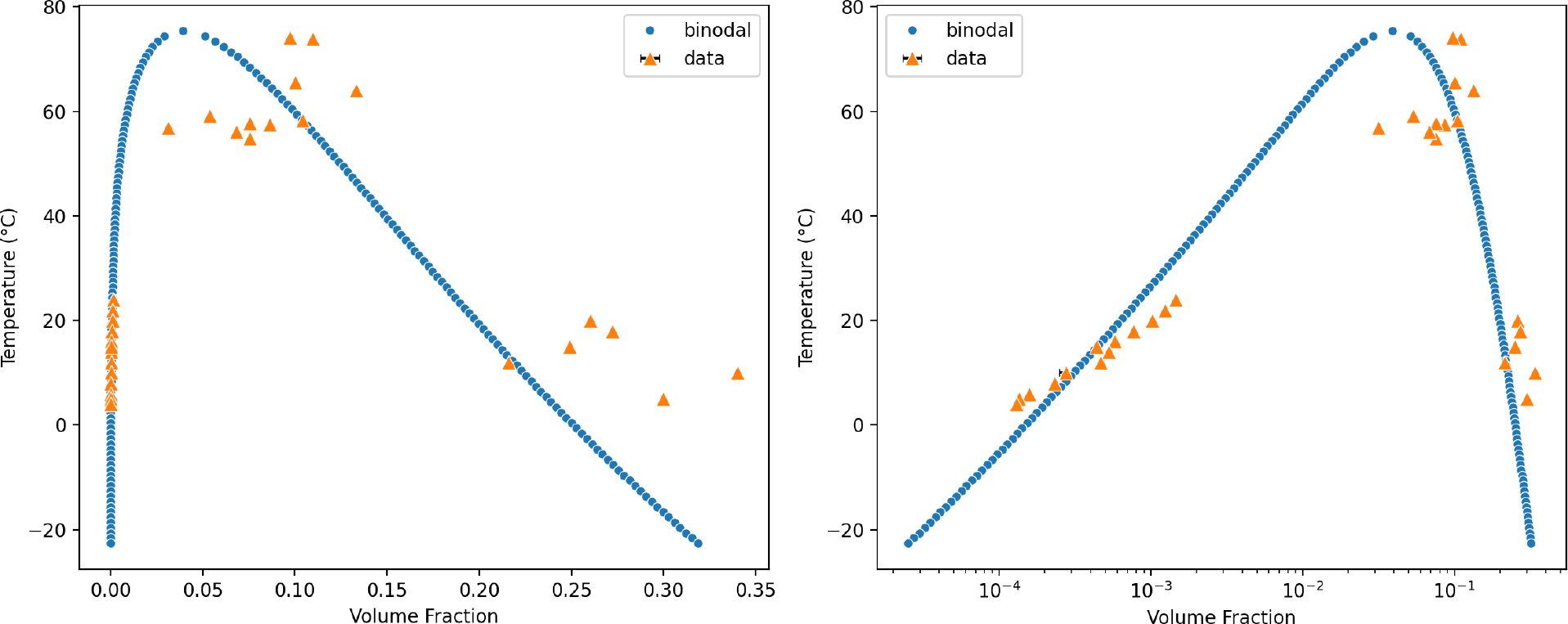
Example output from *fireball-fit* applied to phase separation data of the wild-type A1-LCD from Bremer et al. (48) **using GCT**. This is a four-parameter fit using GCT. Protein concentration is converted to volume fraction by assuming that a volume fraction of 1.0 corresponds to a mass concentration of 1310 mg/mL. The final fitting parameters are θ = 460 K, *w*_2_ = 0.0168, *w*_3_ = 0.00632, and *n* = 1944. The right panel is identical to the left one, save that it uses a logarithmic scale for the abscissa.

*fireball-draw* is used to draw a binodal using a specified free energy model and given parameters. This function may also be used to draw a binodal alongside a dataset, to determine the adequacy of a given set of parameters in describing a known binodal. *fireball-draw* saves the binodal to the disk and outputs the binodal to the screen.

*fireball-single-chain-fit* and *fireball-single-chain-draw* are functional analogs of *fireball-fit* and *fireball-draw* that are used to fit or draw a single-chain coil-to-globule transition curve using GCT to fit data from simulations or experiments. These functions save the coil-to-globule transitions and plot the fit alongside the data. An example of this output is shown in **Fig. 3**. The fitted parameters from *fireball-single-chain-fit* can be used as inputs for *fireball-draw* to predict phase diagrams based on single-chain behaviors.

**Fig. 3:**
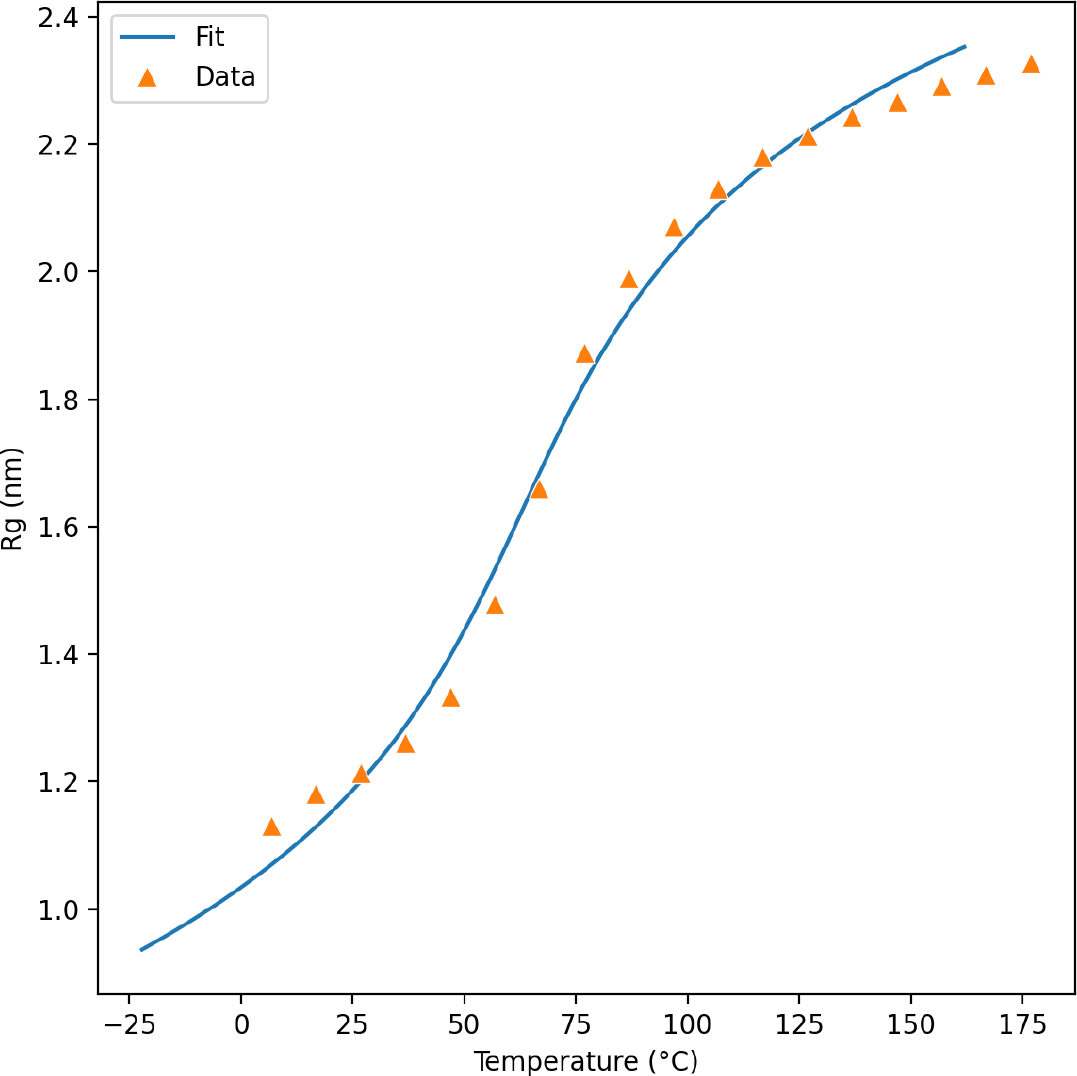
Example output of *fireball-single-chain-fit* applied to all-atom simulation data of the (QGQSPYG)_9_ protein from Zeng et al. (53). This is a four-parameter fit using GCT with a Flory characteristic ratio of 2.5, a Kuhn length of 7, and a monomer length of 0.38 nm. The final fitting parameters are θ = 452 K, *w*_3_ = 0.118, *w*_3_ = 0.00830, and *n* = 92.0. The extrapolation to temperatures below 0°C are shown to generate a smooth curve.

## Discussion

Here we introduce FIREBALL, a computational suite to enable analysis and fitting of data from measurements or computations that provide information regarding the concentrations of coexisting phases as a function of temperature or other control parameters. FIREBALL is designed as an easy-to-use method for generating phase diagrams based on various theoretical models and functional forms for the free energies of polymer solutions. Although support is currently provided for the FHT and GCT models, one can readily include support for other free energy models. FIREBALL is intended for research and pedagogy. It can be used for quick estimations of sequence-specific parameters of phase separation, allowing comparative analyses across different data sets. Importantly, FIREBALL also allows one to determine when mean-field homopolymer theories, such as Flory-Huggins theory and Gaussian cluster theory, break down in describing experimental data.

In addition to phase separation data, FIREBALL also allows one to analyze and fit coil-to-globule transition data. There is now a large corpus of data demonstrating the connection between single-chain conformations and multi-chain phase behaviors of polymers (27,45–50,41). As the theory behind this connection continues to be clarified, the functionality implemented in FIREBALL offers a method for going directly from single-chain data to phase diagrams.

Lastly, we have built FIREBALL with ease-of-use in mind. This approach also extends to adding new functionalities to FIREBALL. Within the documentation, we explain how to add new theoretical models, which only requires one to write a single file that defines the free energy-based functions. The long-term goal with FIREBALL, which we are releasing as an open-source, public resource, is to provide a toolbox that allows others to generate, analyze, and fit phase diagrams and coil-to-globule transitions, while also promoting innovation in the field of biological phase separation.

## Author Contributions

M.F., A.S.H., and R.V.P. conceptualized FIREBALL. M.F. and A.S.H. wrote the software. X.Z. assisted to include Gaussian cluster theory into FIREBALL. M.F. performed the analysis to prototype the software. M.F. and R.V.P. wrote the manuscript.

## Declaration of Interests

R.V.P. is a member of the Scientific Advisor Board of Dewpoint Therapeutics. The work reported here was not influenced by this affiliation. All other authors declare no competing interests.

## Acknowledgments

This work was supported by grants from the US National Institutes of Health (grant R01NS121114) and the Air Force Office of Scientific Research (grant FA9550-20-1-0241).

## Notes

https://github.com/Pappulab/FIREBALL

